# Brain Structure and Working Memory Adaptations Associated with Maturation and Aging in Mice

**DOI:** 10.1101/2023.02.24.529940

**Authors:** Kevan P. Clifford, Amy E. Miles, Thomas D. Prevot, Keith A. Misquitta, Jacob Ellegood, Jason P. Lerch, Etienne Sibille, Yuliya S. Nikolova, Mounira Banasr

## Abstract

**Introduction:** As the population skews toward older age, elucidating mechanisms underlying human brain aging becomes imperative. Structural MRI has facilitated non-invasive investigation of lifespan brain morphology changes, yet this domain remains uncharacterized in rodents despite increasing use as models of disordered human brain aging.

**Methods:** Young (2m, n=10), middle-age (10m, n=10) and old (22m, n=9) mice were utilized for maturational (young vs. middle-age) and aging-related (middle-age vs. old mice) comparisons. Regional brain volume was averaged across hemispheres and reduced to 32 brain regions. Pairwise group differences in regional volume, residualized for total brain volume, and associations between volume and cognitive performance on the Y-maze task were tested. General linear models with total brain volume as a covariate, and logistic regression for sample wide associations were employed respectively, correcting for multiple comparisons. Structural covariance networks were generated using the R package ‘igraph’. Group differences in network centrality (degree), integration (mean distance), and segregation (transitivity, modularity) were tested across network densities (5–40%), using 5,000 (1,000 for degree) permutations with significance criteria of p<0.05 at ≥5 consecutive density thresholds.

**Results:** Widespread significant maturational changes in volume occurred in 18 brain regions, including considerable loss in isocortex regions and increases in brainstem regions and white matter tracts. The aging-related comparison yielded 6 significant changes in brain volume, including further loss in isocortex regions and increases in white matter tracts. No significant volume changes were observed across either comparison for subcortical regions. Additionally, smaller volume of the anterior cingulate area (χ^2^=2.325, pbh=0.044) and larger volume of the hippocampal formation (χ^2^=-2.180, pbh=0.044) were associated with poorer cognitive performance. Maturational network comparisons yielded significant degree changes in 9 regions, but no aging related changes, aligning with network stabilization trends in humans. Maturational decline in modularity occurred (24-29% density), mirroring human trends of decreased segregation in young adulthood, while mean distance and transitivity remained stable.

**Conclusions/Implications:** These findings offer a foundational account of age effects on brain volume, structural brain networks, and cognition in mice, informing future work in facilitating translation between rodent models and human brain aging.

## 1. Introduction

As the world population skews sharply toward older age, with 21.1% projected to be 60 years or older by 2050, the risk of age-related physical and cognitive decline is expected to increase an already widespread societal burden (Chatterji et al. 2015). Understanding lifespan changes in brain maturation and aging is crucial for uncovering etiology driving the aging process and the emergence of age-related diseases, such as dementia (Hou et al. 2019).

Human structural magnetic resonance imaging (MRI) studies have begun to map trajectories of lifelong structural brain change. In humans, overall brain volume is observed to increase before a mid-childhood peak at 5.9 years, followed by a near-linear decrease across the lifespan (Bethlehem et al. 2022). On a regional level, maturational growth peaks earliest in primary sensory motor areas and latest in higher-order association areas (Giedd and Rapoport 2010). Specifically, regional gray matter loss occurs most consistently in the cerebral cortex, with drastic loss in prefrontal regions accompanied by less extensive loss in parietal and temporal association cortices (Resnick et al. 2003; Tisserand et al. 2002; N Raz 1997). Moderately consistent findings of aging-related volume loss are reported for the insula, cerebellum, basal ganglia, and thalamus (Good et al. 2001; G. Alexander et al. 2012). However, findings of aging-related atrophy in limbic and prelimbic regions are more conflicting, particularly in the hippocampus, amygdala, and cingulate gyrus (Jernigan et al., 2001; Raz, 1997; Zheng et al., 2019). Compared to gray matter, white matter volume matures gradually, peaking at 50 years of age, before drastically declining after 60 years of age (Jernigan et al. 2001; Liu et al. 2016). In turn, age-related individual differences in regional gray and white matter properties have been mapped onto differences in cognition and may contribute to observed patterns of age-dependent cognitive decline (Oschwald et al., 2019; Raz & Rodrigue, 2006a).

While structural MRI has facilitated progress in characterizing lifespan brain volume changes in humans, further leveraging these insights to study molecular mechanisms of aging is challenging due to the invasive nature of molecular brain research. The use of rodent models allows for more direct experimentation than human brain research, including transgenic and gene knock out models to simulate pathology, drug trials, and more invasive biological interventions (Nadon 2006). Moreover, the significantly shorter lifecycles in rodent models facilitate feasible lifespan and generational research (Folgue ras et al. 2018). Ultimately, models are as useful as their ability to approximate healthy and pathological states in the human brain. However, while preclinical models are being increasingly implemented for Alzheimer’s disease, Parkinson’s disease, frontotemporal dementia, and Huntington’s disease (Ellenbroek and Youn 2016; Dawson, Golde, and Lagier-Tourenne 2018), normal lifespan changes in brain volume remain incompletely characterized in rodent models.

This incomplete characterization reflects, in part, evolving methodology such as specialized small-rodent scanners and non-standardized brain segmentation methods (Sawiak et al. 2013; Denic et al. 2011), that has contributed to a lag in the use of structural MRI in preclinical rodent models compared to its widespread use in human research (Feo and Giove 2019). As a result, there are few lifespan brain volume studies in rodents. Whole-brain volumetric findings suggest that gray matter volume increases in rodents before plateauing at 2 months (Mengler et al. 2014). A recent study describes prominent regional volume loss in the isocortex (i.e., often referred to as neocortex in humans), most notably in the prefrontal cortex and temporal association areas, as well as the insula and perirhinal area. Further loss occurred in the cerebellum, while increases in volume occurred in somatosensory areas, thalamus, midbrain, septal nuclei, and specific hippocampal regions (G. E. Alexander et al. 2020a). The sparse literature on lifespan brain volume changes in rodents poses a gap in the determination of their suitability as a model of healthy structural brain aging.

In this context, specific brain regions warrant particular attention due to their selective age sensitivity and relevance to age-related cognitive phenotypes. These regions of interest (ROIs) include the anterior cingulate cortex (ACC), hippocampus, and orbitofrontal cortex (OFC). In humans, the ACC connects to both prefrontal and limbic regions, integrating emotional and cognitive functions, including attention, decision making, fear response, and memory (Bush, Luu, and Posner 2000). In rodents, the homologous anterior cingulate area subserves similar functions to those of the ACC (Zhang et al. 2014; Hosking, Cocker, and Winstanley 2014; Frankland et al. 2004), comprising one of four subregions of the medial prefrontal cortex (Heidbreder and Groenewegen 2003). Normal aging is largely associated with volume loss in the ACC (Bergfield et al. 2010; Good et al. 2001), although preservation has also been reported (N Raz 1997). Further evidence of aging-dependent changes in the ACC includes reductions in spine density and dendritic tree extent (Markham and Juraska 2002), and decreases in glucose uptake and metabolism (Pardo et al. 2007).

The hippocampus is another region of particular relevance due to its sensitivity to aging related changes. By integrating sensory, spatial, and temporal information, it contributes to memory formation, learning, and spatial navigation (Eichenbaum 1999; Maguire, Burgess, and O’Keefe 1999). In humans, drastic hippocampal volume loss is observed in aging (Jernigan et al. 2001; Zheng et al. 2019), however conflicting results have also been reported (Good et al. 2001; E. Sullivan 1995). Furthermore, this region displays a unique pattern of continual neurogenesis and increase in brain volume into adulthood in both humans and rodents (E. Sullivan 1995; E. V. Sullivan et al. 2006; Diamond, Johnson, and Ingham 1975), while reduced hippocampal neurogenesis in late life has been associated with decreased volume and memory performance (Bettio, Rajendran, and Gil-Mohapel 2017; Driscoll et al. 2006; Kuhn, Dickinson-Anson, and Gage 1996).

Lastly, the OFC in humans is a prefrontal cortical region that acts as a link between sensory integration, autonomic reactions, learning, and decision making for emotional and reward-related behaviours (Kringelbach 2005). This age-sensitive region is of interest due to evidence of decreasing volume across aging (Tisserand et al. 2002; Convit et al. 2001). In rodents, the orbital area is homologous to a human agranular section of the OFC, serving similar function (Öngür and Price 2000; Izquierdo 2017). Better understanding the nature of age-dependent structural changes in these regions is likely to provide insight into age-dependent cognitive changes.

Network-based approaches to modeling brain structure can complement univariate analyses of regional brain volume to uncover mechanisms underlying normal aging. Structural covariance networks (SCNs) are comprised of nodes (brain regions) connected by edges which represent the statistical correlation between properties of adjacent nodes (Bullmore and Sporns 2009). As such, they reflect synchronized changes in morphology (volume, cortical thickness) between brain regions (Alexander-Bloch, Giedd, and Bullmore 2013), thought to be driven by mutually trophic influences (Ferrer et al. 1995; Pezawas et al. 2004) and experience-driven plasticity (Mechelli 2005). SCNs are heritable (Mechelli 2005), occur at the individual and population level (Alexander-Bloch, Giedd, and Bullmore 2013; Tijms et al. 2012), and partially map on to structural and functional connectivity (Alexander-Bloch, Giedd, and Bullmore 2013). SCNs also display consistent lifespan trajectories, reflecting coordinated neurodevelopmental changes across maturation and aging (Fan et al. 2011; Khundrakpam et al. 2013; Cao et al. 2016). Furthermore, perturbations to typical composition of SCNs have been observed in neurodegenerative diseases and psychiatric illnesses (Alexander-Bloch, Giedd, and Bullmore 2013; Prasad et al. 2022).

While SCN analysis could provide additional insight into age-related changes in brain structure and organization, few studies have considered lifespan SCN trajectories in rodents. Existing literature has employed scaled subprofile modelling SCN methods (G. E. Alexander et al. 2020b), but none, to our knowledge, have leveraged whole-brain SCN and graph theoretical methods. The latter can be used to characterize different aspects of SCN topology, from which inferences regarding brain structure and function can be drawn (Bullmore and Sporns 2009). Topological measures broadly index network (1) node centrality, identifying the ‘hubness’ of highly connected and influential nodes; (2) integration, capturing the capacity for efficient network-wide communication and information processing; and (3) segregation, quantifying the presence of densely interconnected and clustered nodes facilitating local or specialized information processing (Bullmore and Sporns 2009; Rubinov and Sporns 2010).

In the present study, we aim to use structural MRI and graph theoretical methods to provide a comprehensive characterization of lifespan brain volume changes in mice. By comparing young mice to middle-aged mice, and middle-aged mice to old mice, we distinguish between maturational and aging-related differences, respectively, in regional brain volume for 32 brain regions. Particular consideration is given to the three aforementioned *a priori* ROIs, for which there is evidence of age-sensitivity and relevance to age-related cognitive phenotypes (anterior cingulate area, hippocampal formation, and orbital area). We then map these regional brain volume changes to cognitive outcomes by establishing associations with working memory performance on the Y maze task. Lastly, we further explore regional volumetric changes by analyzing maturational and aging-related differences in SCNs, examining network topology across the mouse lifespan.

## 2. Methods

### 2.1 Animals

Two months-old (young), 10 months-old (middle-age) and 22 months-old (old) male C57BL/6 mice (n=10/group, Charles River Laboratories, Senneville, Quebec, Canada) were housed under a 12/12-hours light/dark cycle (7am-7pm light phase) at constant temperature (20-23°C) with free access to food and water. One mouse in the 22 months-old animal group was excluded from the study for developing cataract signs before behavioral testing. Testing was performed by an experimenter blinded to the animal group during the animal’s light cycle phase. Animal use and testing procedures were conducted in accordance with the Canadian Council for Animal Care with approval from the institutional animal care committee.

### 2.2. Behavioral testing

#### Y-maze

*Testing was performed as in Prevot et al 2018, 2021.* The Y-maze was chosen in preference over other working memory tasks (e.g., Morris water maze or radial arm maze) as it is relatively less susceptible to confounding factors related to aging, such as weight, fatigue, and mobility differences (Matzel et al., 2008). The Y-maze apparatus consists of 3 identical black PVC arms (26 cm length x 8 cm width x 13 cm height) in the shape of a Y; each arm has a sliding door. Distal cues of different shape, form and color were placed on the walls around the room. Mice were allowed to freely explore the apparatus for 10 minutes on 2 consecutive days (*habituation stage*). The next day, mice performed a training session consisting of seven consecutive trials. Each trial began with placing the animal in the starting arm for 5 seconds, prior to the door of y-maze arm opening, allowing the animal to freely choose between the 2 goal arms, and choices were recorded. Upon entering the arm, the door was closed for a 30s period (inter-trial interval or ITI) after which the animal was returned to the starting arm for the next trial (*training stage*). The following day, mice were subjected to a similar procedure as the training session, with the exception that the ITI being lengthened to 60s ITI *(testing stage)*. The mean percent alternation rate (number of alternation/number of trials × 100) was calculated as an index of working memory.ANOVA was performed to determine differences between groups, followed by the Wald test in a logistic regression model, to account for the non-normality of the data distribution and the non-continuous nature of the proportion alternation scores.

#### Locomotor activity

During the light cycle, locomotor activity of each mouse was measured for 1 hour in an observational cage (30 x 30cm) with bedding on the floor (Noldus Phenotyper®, Leesburg, VA, USA). Using NoldusEthoVision 10 tracking software, mice total distance traveled within the cage was analyzed.

### 2.3 MRI data collection and preprocessing

Twenty-four hours after the last behavioral testing, animals were anesthetized and perfused with 4% paraformaldehyde containing 2mM of ProHance (gadoteridol). Brains were then prepared and scanning was performed as described in (Nikolova et al. 2018). In this case, however, we extracted brain volumes from a total of 280 regions (instead of 159, as in Wheeler et al. 2015), excluding ventricles. This parcellation method was adapted from previous rodent MRI studies (Dorr et al. 2008; Richards et al. 2011; Steadman et al. 2014; Ullmann et al. 2013). Deformation-based morphometry was used to derive absolute brain volumes (in mm^3^). For all MRI analyses, ROI volumes were averaged across hemispheres in order to minimize multiple testing burden.

### 2.4 Data reduction

MRI-derived volumes were extracted from 280 regions. To reduce the number of comparisons, these regions were combined according to their positions within a hierarchical tree, ultimately yielding 32 regions of interest (ROIs) (Refer to Supplementary Material 1). Of these ROIs, 5 are white matter tracts (cerebellar-related fiber tracts, cranial nerves, extrapyramidal fiber system, lateral forebrain bundle system, medial forebrain bundle system) and 27 are gray matter divisions. The latter includes 5 brainstem areas (hypothalamus, medulla, midbrain, pons, thalamus); 2 cerebral nuclei (pallidum, striatum); 18 cerebral cortical areas, including the cortical subplate, hippocampal formation, olfactory areas, and 15 isocortical areas (agranular insular area, anterior cingulate area, auditory areas, ectorhinal area, frontal pole, infralimbic area, orbital area, perirhinal area, posterior parietal association areas, prelimbic area, retrosplenial area, somatomotor areas, somatosensory areas, temporal association areas, visual areas); and 2 cerebellar divisions (cerebellar cortex, cerebellar nuclei).

### 2.5 Volumetric Analyses

#### 2.5.1 *A priori* ROI volumetric effects of age

Preliminary statistical analyses focused on the anterior cingulate area, hippocampal formation, and orbital area. For each ROI, we performed the following analyses. First, we tested pairwise group differences (young vs. middle-age, middle-age vs. old) in volume using separate general linear models including total brain volume as a covariate. Then, we tested the sample-wide associations of regional volume, residualized for total brain volume, with working memory performance. Working memory performance was evaluated using the Y maze task, and logistic regression was employed with volume measurement as a continuous independent variable and percent alteration expressed as a proportion score, bound between the values of 0 and 1, as the dependent variable (J. S. Long, 1997). To account for multiple comparisons, raw p-values from each analysis (n=6, 3, respectively) were adjusted using the Bonferroni-Holm method (Holm, 1979), a correction commonly used in MRI analysis (Kerr et al., 2022; Straub et al., 2023). We opted for this relatively conservative correction method to ensure greater specificity of our findings. Notably, however, the Bonferroni-Holm method is less conservative than the Bonferroni correction, offering similar protection against Type I error coupled with reduced likelihood of Type II error (Eichstaedt et al., 2013).

#### 2.5.2 Whole-brain volumetric effects of age

We repeated the aforementioned analyses brain-wide, correcting for multiple testing in all 32 ROIs using the Bonferroni-Holm method.

### 2.6 Structural covariance network analyses

#### 2.6.1 *A priori* degree centrality analyses

We used the R package ‘igraph’ to generate and compare age group-specific structural covariance matrices indexing correlations of volume between brain regions. Regional volumes were first adjusted for total brain volume, and all negative correlations were removed (to align with previous work (Misquitta et al., 2021; Nikolova et al., 2018)), resulting in group-specific structural covariance matrices that were unweighted and unsigned. For each group, a set of structural covariance networks was then defined by thresholding the covariance matrices using a range of density thresholds (0.05-0.40), indexing the top 5-40% strongest connections in sequential increments of 1%. We then tested pairwise group differences in degree centrality, a node’s level of interconnectedness within the network (Bullmore and Sporns, 2009), for each of the 3 *a priori* ROIs (anterior cingulate area, hippocampal formation, and orbital area) across each of the above mentioned age group-specific structural covariance networks. For statistical analysis, permutation testing was performed at each density threshold (n=5000 permutations), yielding null distributions against which empirical two-tailed p-values were computed. Significance was defined as p<0.05 at 5 or more consecutive density thresholds.

#### 2.6.2 Brain-wide degree centrality analyses

We repeated the degree centrality analyses for all 32 brain regions. Permutation testing was performed at density thresholds using n=1000 permutations.

#### 2.6.3 Global structural covariance network analyses

In addition, we conducted pairwise group comparisons (young vs. middle-age, middle-age vs. old) of modularity and transitivity, network segregation measures indexing densely internally connected subgraphs within the network, and the tendency of nodes to form clusters, respectively; and mean distance, the average length of shortest paths between nodes, reflecting overall network efficiency (Bullmore and Sporns, 2009; Rubinov and Sporns, 2010). Permutation testing was performed across a range of density thresholds (0.05-0.40), using n=5000 permutations. To further explore group differences in network modularity, a Walktrap community detection algorithm from the ‘igraph’ package was used to identify densely connected subgraphs within age-group specific networks. For each age group, the most prominent provincial hubs and connector hubs were identified in communities by calculating both the within-community degree z-score (the number of connections a given node has to other nodes within the same community) and the participation coefficient (the distribution of a node’s connections to other communities in the network) of nodes, as outlined in (Guimerà et al. 2005) and adapted from (Shizuka 2019). Provincial hubs drive exchange and integration of information within a single segregated community. As such, they are defined by a high within-community degree z-score and a low participation coefficient. In contrast, connector hubs facilitate sharing and integration of information between otherwise segregated communities and are therefore defined by both a high within-community degree z-score and a high participation coefficient (Sporns 2016).

## 3. Results

### 3.1 Cognitive performance on Y-maze task and brain volume

Although old mice showed average percent alternations close to 50% chance compared to that of 70% in young animals, we found no significant difference in the percent of spontaneous alternation between young, middle-age, or old animal groups (ANOVA F_2,26_=1.96, p=0.16, Figure 1A). Follow up analysis using the Wald test in a logistic regression model similarly found no significant difference in proportion of alternation between age groups (χ^2^=4.7, p=0.095). We did not detect any statistical difference between groups in distance traveled in the locomotor activity test (F_1,26_=2.240, p=0.117; Young: 6311cm±133cm; Middle-age: 8437cm±795cm; Old: 7215cm ±1141cm). In the absence of between-group differences in cognitive performance, we then examined potential associations between working memory measured in the Y-maze test and brain volume of our 3 *a priori* ROIs and brain-wide (i.e., across all 32 regions) in the full sample. Independent of age group, better cognitive performance was associated with greater volume of the anterior cingulate area (χ^2^=2.325, pbh=0.044, Figure 1B) and smaller volume of the hippocampal formation (χ^2^=-2.180, pbh=0.044, Figure 1C), while no association emerged with volume of the orbital area (χ^2^=0.855, pbh=0.393). Significant brain-behavior associations are depicted in Figure 1, and corresponding test statistics are summarized in Table 1. At the whole-brain level, no association between Y-maze performance and regional volume survived correction for multiple comparisons. However, we identified several associations, wherein a “younger” brain phenotype was associated with better cognitive performance, regardless of chronological age (i.e., across age groups). Specifically, better cognitive performance was associated with larger volume of the agranular insular area (χ^2^=2.256, p=0.024) and ectorhinal area (χ^2^=2.646, p=0.008), both of which were larger in younger relative to middle-aged mice. Conversely, better cognitive performance was also associated with lower volume of the medial forebrain bundle system (χ^2^=-1.968, p=0.049), which was larger in older age.

**Fig. 1:**
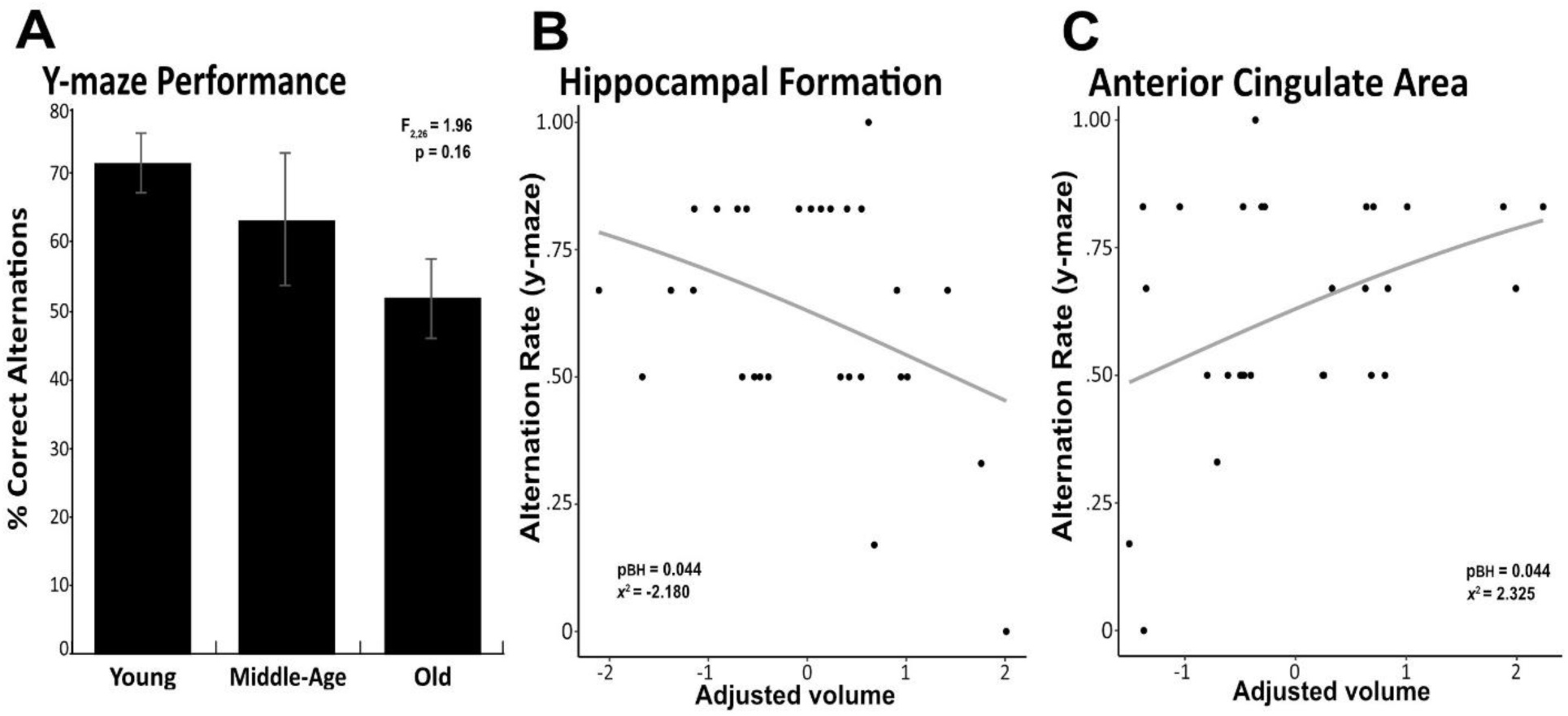
Significant associations between brain volume and working memory performance. Proportion of correct alterations in the Y-maze task are depicted for the young, middle-age and old mice, with error bars representing standard error of the mean (SEM) (**A**). Greater volume of the anterior cingulate area was associated with better working memory performance on the Y-maze task (**B**). Smaller volume of the hippocampal formation was associated with better working memory performance on the Y-maze task **(C).** Associations survived correction for multiple comparisons across these *a priori* regions (p-value pictured) but not across brain-wide regions.

**Table 1.** Sample-wide associations between volume and working memory performance.

| | $\chi^2$ | p<sub>BH</sub> |
| --- | --- | --- |
| **Apriori regions** |  |  |
| Hippocampal Formation | **-2.180** | **0.044** |
| Anterior Cingulate Area | **2.325** | **0.044** |
| Orbital Area | 0.855 | 0.393 |
| **Brain-wide regions** |  |  |
| Brainstem | -- | -- |
| Hypothalamus | -0.517 | 0.645 |
| Medulla | -1.835 | 0.243 |
| Midbrain | -1.822 | 0.243 |
| Pons | -1.475 | 0.299 |
| Thalamus | 0.603 | 0.616 |
| Cerebellum | -- | -- |
| Cerebellar Cortex | 1.418 | 0.312 |
| Cerebellar Nuclei | -1.168 | 0.616 |
| Cerebral Cortex | -- | -- |
| Cortical Subplate | 0.599 | 0.616 |
| Hippocampal Formation | -2.180 | 0.234 |
| Olfactory Areas | 0.885 | 0.523 |
| Cerebral Isocortex | -- | -- |
| Agranular Insular Area | 2.256 | 0.234 |
| Anterior Cingulate Area | 2.235 | 0.234 |
| Auditory Areas | 1.348 | 0.316 |
| Ectorhinal Area | 2.646 | 0.234 |
| Frontal Pole | -0.586 | 0.616 |
| Infralimbic Area | 0.466 | 0.662 |
| Orbital Area | 0.855 | 0.523 |
| Perirhinal Area | 1.243 | 0.342 |
| Posterior Parietal Association Areas | 0.184 | 0.854 |
| Prelimbic Area | 0.617 | 0.616 |
| Retrosplenial Area | 0.867 | 0.523 |
| Somatomotor Areas | 1.359 | 0.316 |
| Somatosensory Areas | 1.311 | 0.320 |
| Temporal Association Areas | 1.670 | 0.302 |
| Visual Areas | 0.608 | 0.616 |
| Cerebral Nuclei | -- | -- |
| Pallidum | -1.418 | 0.299 |
| Striatum | 1.837 | 0.243 |
| White Matter Tracts | -- | -- |
| Cerebellum Related Fiber Tracts | -1.625 | 0.278 |
| Cranial Nerves | -1.532 | 0.299 |
| Extrapyramidal Fiber System | -1.163 | 0.373 |
| Lateral Forebrain Bundle System | -1.941 | 0.243 |
| Medial Forebrain Bundle System | -1.968 | 0.243 |

Sample-wide associations between regional volume, residualized for total brain volume, and working memory performance, indexed as percent correct entries on the Y-maze task. *BH* = Bonferroni-Holm

### 3.2 Volumetric changes across the lifespan

#### 3.2.1 *A priori* ROI volumetric effects of age

We detected significant main effects of age on regional volume of the anterior cingulate area and orbital area, but not of the hippocampal formation. Specifically, we found a maturational decrease in volume in the anterior cingulate area, which was smaller in middle-aged mice than in young mice (t=-4.898, pbh=0.001) (Figure 2A), but did not differ between middle-aged and old mice. We observed a different trajectory of volume loss in the orbital area, where volume was significantly lower in old mice than in middle-aged mice (t=-2.859, pbh=0.038; Figure 2B), but did not differ between young and middle-aged mice, indicating aging-related volume loss. Results of all pairwise comparisons are summarized in Table 2.

**Fig. 2:**
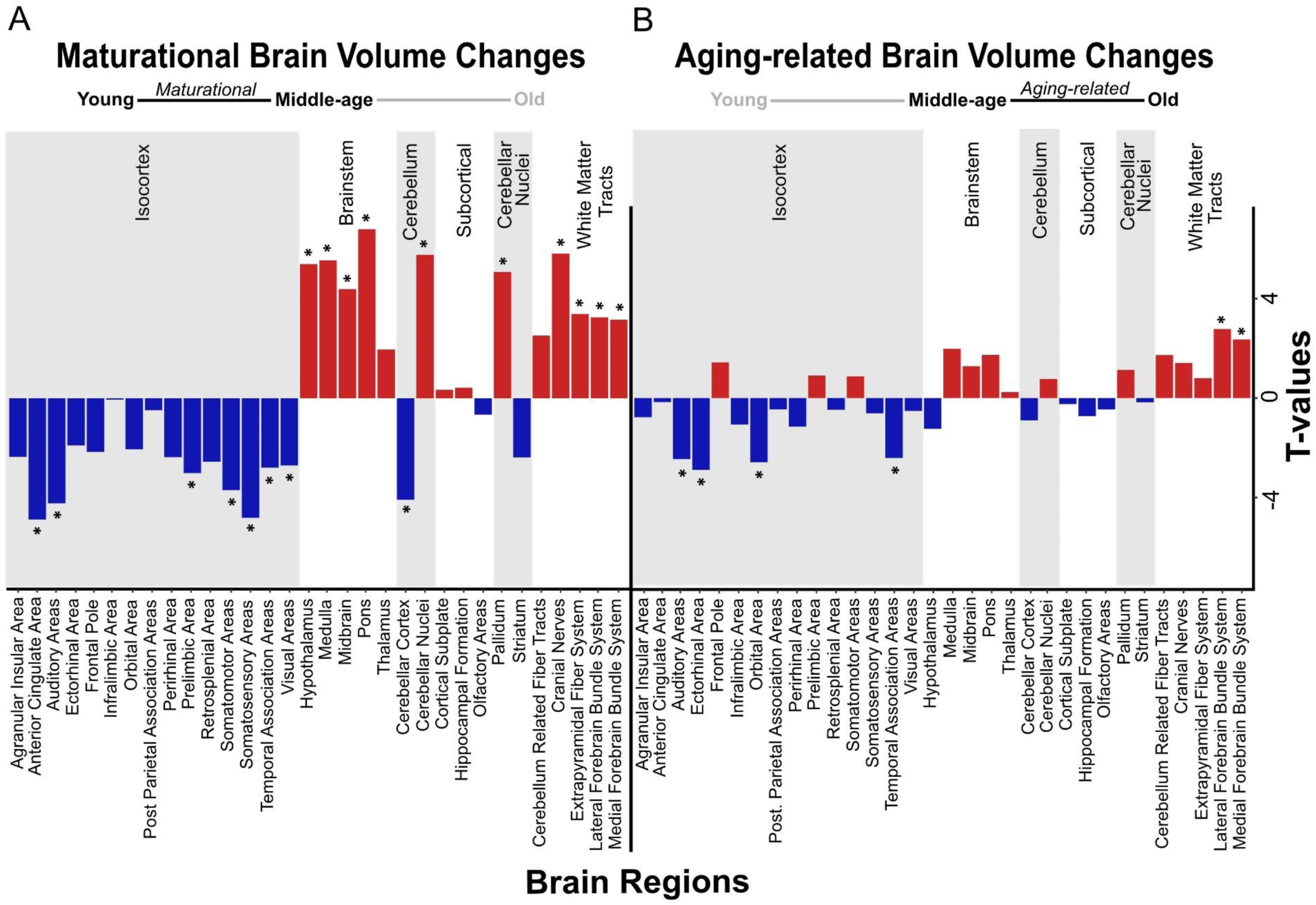
Volumetric changes in maturational and aging-related comparisons. Negative and positive linear changes in regional brain volume are depicted between pairwise comparisons of young and middle-age mice (maturational) (**A**) and middle-age and old mice (aging-related) (**B**). Brain regions that underwent significant change in volume are denoted by *

**Table 2.** Pairwise group differences in volume.

|  | Young vs. Middle-age |  | Middle-age vs. Old |  |
| --- | --- | --- | --- | --- |
|  | t | pBH | t | pBH |
| **Apriori regions** |  |  |  |  |
| Hippocampal Formation | 0.428 | 0.744 | -0.810 | 0.550 |
| Anterior Cingulate Area | **-4.898** | **0.001** | -0.181 | 0.872 |
| Orbital Area | -2.060 | 0.111 | **-2.859** | **0.038** |
| **Brain-wide regions** |  |  |  |  |
| Brainstem | -- | -- | -- | -- |
| Hypothalamus | **5.425** | **0.001** | -1.374 | 0.301 |
| Medulla | **5.581** | **0.001** | 2.197 | 0.092 |
| Midbrain | **4.413** | **0.003** | 1.419 | 0.287 |
| Pons | **6.837** | **0.000** | 1.923 | 0.132 |
| Thalamus | 1.975 | 0.126 | 0.268 | 0.831 |
| Cerebellum | -- | -- | -- | -- |
| Cerebellar Cortex | **-0.410** | **0.004** | -0.996 | 0.475 |
| Cerebellar Nuclei | **5.802** | **0.001** | 0.852 | 0.531 |
| Cerebral Cortex | -- | -- | -- | -- |
| Cortical Subplate | 0.347 | 0.795 | -0.272 | 0.831 |
| Hippocampal Formation | 0.428 | 0.744 | -0.810 | 0.550 |
| Olfactory Areas | -0.658 | 0.640 | -0.509 | 0.710 |
| Cerebral Isocortex | -- | -- | -- | -- |
| Agranular Insular Area | -2.365 | 0.067 | -0.857 | 0.531 |
| Anterior Cingulate Area | **-4.898** | **0.001** | -0.181 | 0.872 |
| Auditory Areas | **-4.243** | **0.004** | **-2.725** | **0.044** |
| Ectorhinal Area | -1.901 | 0.132 | **-3.199** | **0.022** |
| Frontal Pole | -2.160 | 0.094 | 1.586 | 0.229 |
| Infralimbic Area | -0.048 | 0.962 | -1.180 | 0.380 |
| Orbital Area | -2.060 | 0.111 | **-2.859** | **0.038** |
| Perirhinal Area | -2.377 | 0.067 | -1.277 | 0.343 |
| Posterior Parietal Association Areas | -0.478 | 0.717 | -0.504 | 0.710 |
| Prelimbic Area | **-3.021** | **0.028** | 1.000 | 0.475 |
| Retrosplenial Area | -2.565 | 0.051 | -0.522 | 0.710 |
| Somatomotor Areas | **-3.711** | **0.009** | 0.962 | 0.487 |
| Somatosensory Areas | **-4.827** | **0.001** | -0.681 | 0.634 |
| Temporal Association Areas | **-2.799** | **0.039** | **-2.674** | **0.047** |
| Visual Areas | **-2.711** | **0.044** | -0.575 | 0.692 |
| Cerebral Nuclei | -- | -- | -- | -- |
| Pallidum | **5.104** | **0.001** | 1.253 | 0.348 |
| Striatum | -2.384 | 0.067 | -0.190 | 0.872 |
| White Matter Tracts | -- | -- | -- | -- |
| Cerebellum Related Fiber Tracts | 2.537 | 0.052 | 1.917 | 0.132 |
| Cranial Nerves | **5.851** | **0.001** | 1.570 | 0.231 |
| Extrapyramidal Fiber System | **3.408** | **0.016** | 0.888 | 0.528 |
| Lateral Forebrain Bundle System | **3.275** | **0.020** | **3.075** | **0.027** |
| Medial Forebrain Bundle System | **3.177** | **0.022** | **2.611** | **0.050** |

Group differences in regional volume, residualized for total brain volume. A negative t score indicates a period-specific decrease in volume, and a positive t score indicates a period-specific increase in volume. *BH=Bonferroni-Holm*

#### 3.2.2 Whole-brain volumetric effects of age

We found significant group differences in regional volume of 24 of 32 regions tested, including some of the *a priori* ROIs, as mentioned above (Figure 2A, B). In most of these regions (18/24), volume differed significantly between young mice and middle-aged mice, indicating a maturational effect. The maturational effects were characterized by increases in volume (i.e., middle-aged>young) and decreases in volume (i.e., middle-aged<young) in a roughly equivalent number of regions. Maturational increases in volume (t=2.611–6.837, pbh≤0.050) were observed in ten regions, including the cerebellar nuclei, pallidum, most of the brainstem areas (hypothalamus, medulla, midbrain, pons), and most of the white matter tracts (cerebellum-related fiber tracts, cranial nerves, lateral forebrain bundle system, medial forebrain bundle system) (Figure 2A). Maturational decreases in volume (t=2.674 – 4.898, pbh≤0.050) were observed in 10 regions, including the cerebellar cortex and 7 of 15 isocortical areas: anterior cingulate area, auditory areas, prelimbic area, somatosensory areas, somatomotor areas, temporal association areas, visual areas (Figure 2A).

Aging-related (i.e., middle-aged<old) increases in volume were observed in the lateral forebrain bundle system (t=3.275, pbh=0.020) and medial forebrain bundle system (t=3.177, pbh=0.022), white matter tracts in which we also found maturational increases in volume. Aging-related decreases (i.e., middle-aged>old) in volume were observed in two distinct isocortical areas, the ectorhinal cortex (t=-3.199, pbh=0.022) and the orbital area (t=-2.859, pbh=0.038) (Figure 2B). Results of all pairwise comparisons are summarized in Table 2.

### 3.3 Structural covariance network adaptations

#### 3.3.1 *A priori* ROI degree centrality analyses

Among the 3 *a priori* ROIs, we found significant maturational group differences in regional degree centrality of the anterior cingulate area and hippocampal formation, but not of the orbital area (Figure 3A-C). Degree centrality in the anterior cingulate area was significantly lower in middle aged mice than in young mice (23-40% density) indicating a maturational decrease in regional connectivity (Figure 3A, Figure 4A-C). Conversely, degree centrality in the hippocampal formation was significantly higher in middle-aged mice than in young mice (8-16% density) indicating a maturational increase in regional connectivity (Figure 3B, Figure 4D-F). In contrast, no significant degree centrality differences in *a priori* regions were detected in the aging-related comparisons (Figure 3D-F). Results of all pairwise comparisons are summarized in Supplementary Table 1.

**Fig. 3:**
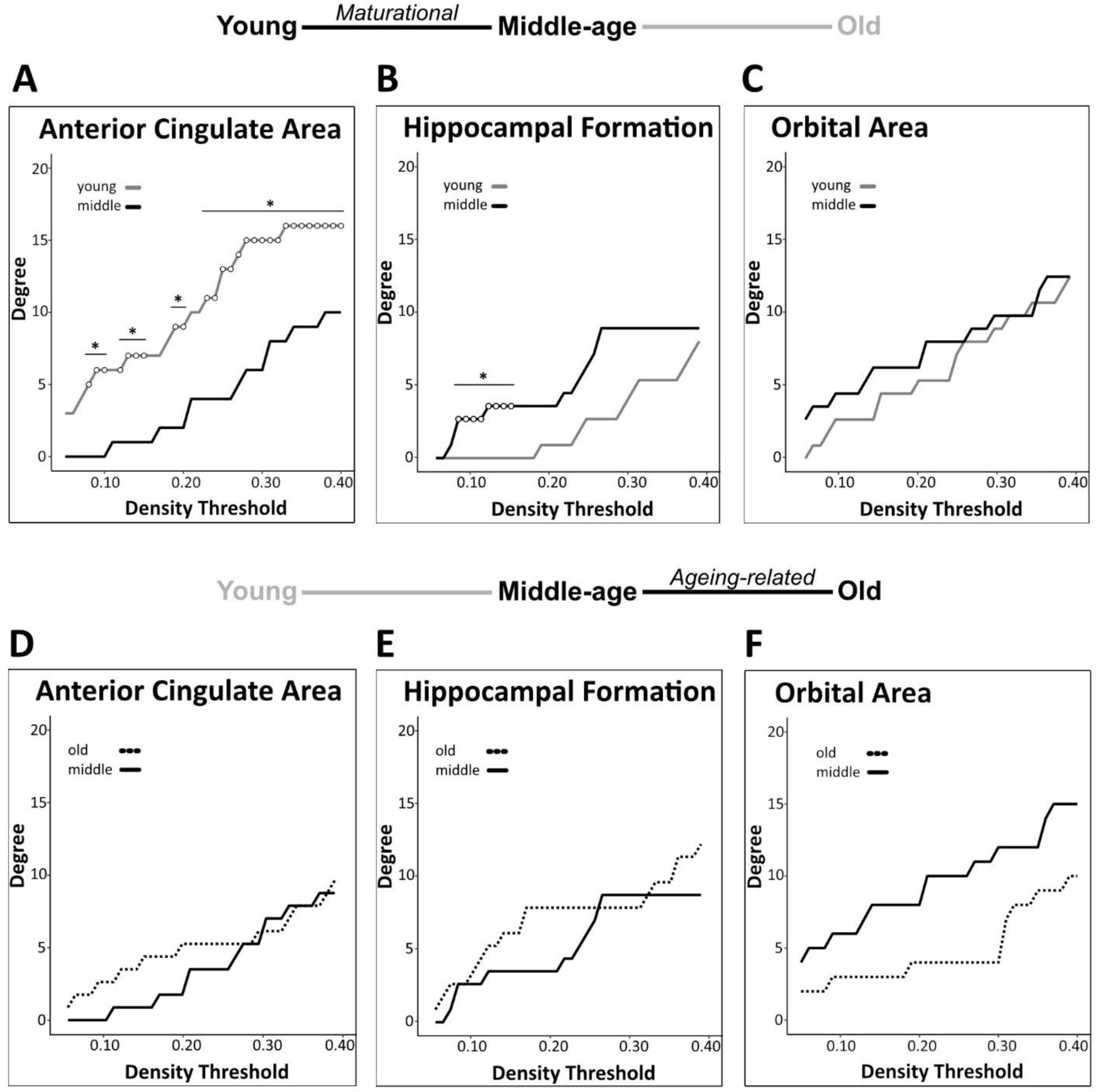
Regional degree centrality differences in *a priori* ROIs for maturational and aging-related comparisons. Maturational comparisons in structural covariance networks for the *a priori* age sensitive regions revealed significant decrease in degree centrality for the anterior cingulate area (**A**), a maturational increase in degree centrality in the hippocampal formation (**B**), and no significant differences observed in the orbital area (**C**). Aging-related comparisons in structural covariance networks for the *a priori* age-sensitive regions did not yield significant differences in degree (**D-F**). Differences in degree were determined across density thresholds of 0.05 and 0.40 at 5 consecutive significant density thresholds (statistical significance indicated by white circles (*p<0.05)).

**Fig. 4:**
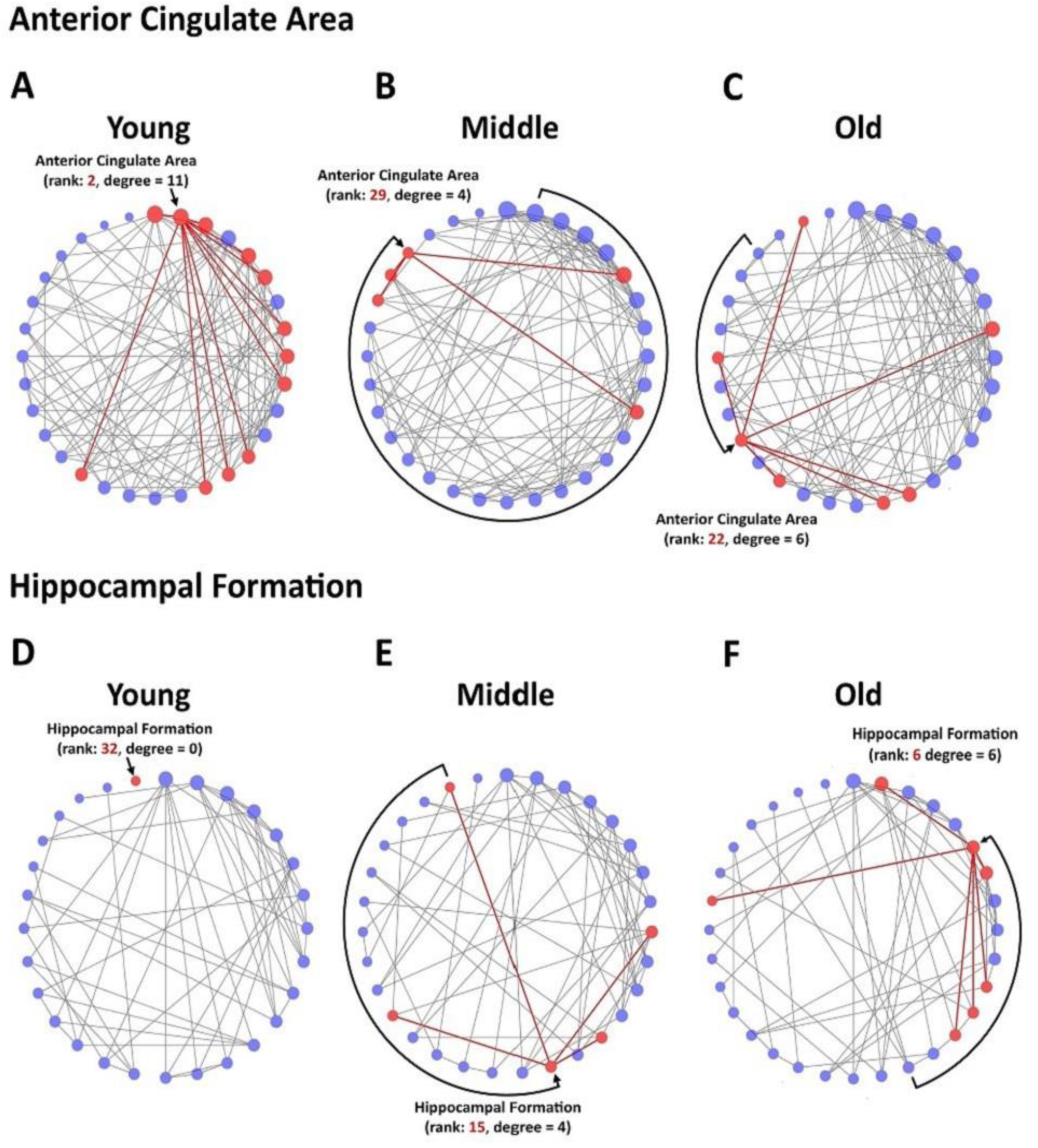
Circular degree plots of significant differences in regional connectivity in *a priori* ROIs. The anterior cingulate area networks in young (**A**), middle-age (**B**) and old (**C**) mice are depicted at network density of 24% for visualization. The hippocampal formation networks in young (**D**), middle-age (**E**) and old (**F**) mice are depicted at network density of 12% for visualization. Nodes are arranged clockwise by degree, the number of significant correlations a given brain region has with other regions within the network, with the highest-degree node at the top of the circle. Degree is further reflected in node size. Rank refers to the numerical value of a region’s degree, sorted largest to smallest in the network. The region of interest and its direct connections are highlighted in red.

#### 3.3.2 Brain-wide degree centrality analyses

We identified group differences in degree centrality of 9 of 32 regions tested. In all of these regions, degree differed significantly between young mice and middle-aged mice, indicating maturational network connectivity changes. In addition to the above-mentioned maturational differences in degree centrality of the anterior cingulate area (decrease) and hippocampal formation (increase), we also detected significant maturational decreases in 3 additional isocortical regions (auditory areas, ectorhinal area, somatomotor areas) and significant maturational increases in 4 regions (cerebellar nuclei, cerebellum-related fiber tracts, cortical subplate, midbrain) across partially overlapping density thresholds (Figure 5A-G). No significant differences in regional degree were found between the middle-aged and old mice.

**Fig. 5:**
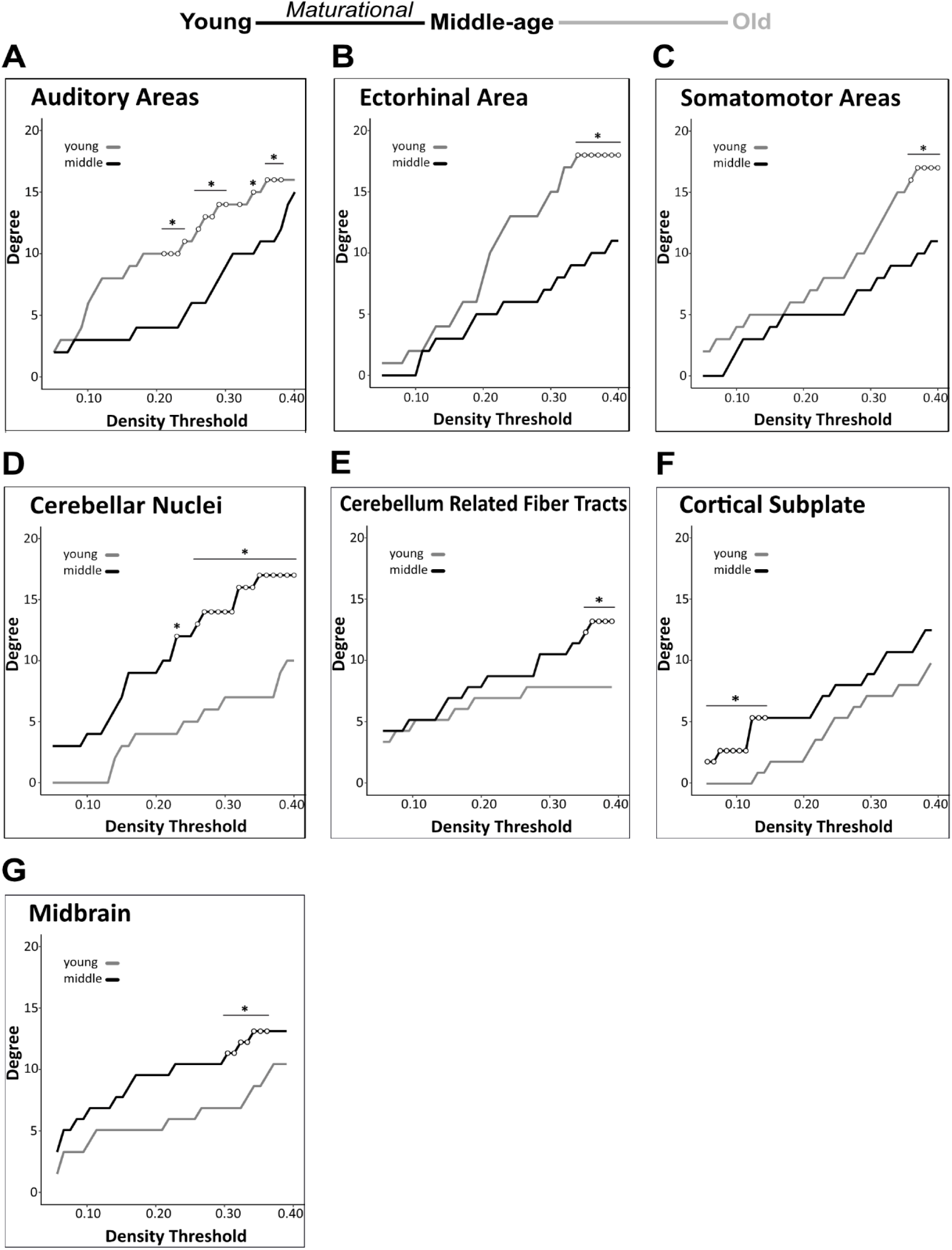
Brain-wide degree centrality differences in maturational and aging-related comparisons. Brain-wide maturational comparisons in structural covariance networks revealed significant decreases in degree in the auditory areas (**A**), ectorhinal area (**B**) and somatomotor areas (**C**), and maturational increases in degree in the cerebellar nuclei (**D**), cerebellum related fiber tracts (**E**), cortical subplate (**F**) and midbrain (**G**). No significant differences were found in any brain regions in the aging-related network comparisons. Changes in degree were determined across density thresholds of 0.05 to 0.40 at 5 consecutive significant density thresholds (statistical significance indicated by white circles (*p<0.05)).

#### 3.3.3 Global structural covariance network analyses

We detected a significant maturational decrease (i.e., middle-age<young) in global network modularity (density 24-29%) (Figure 6A), but no significant aging-related differences therein (Figure 6B). In addition, we did not detect any significant maturational or aging-related differences in global network transitivity or mean distance measures. Results of all pairwise comparisons are summarized in Supplementary Table 2.

**Fig. 6:**
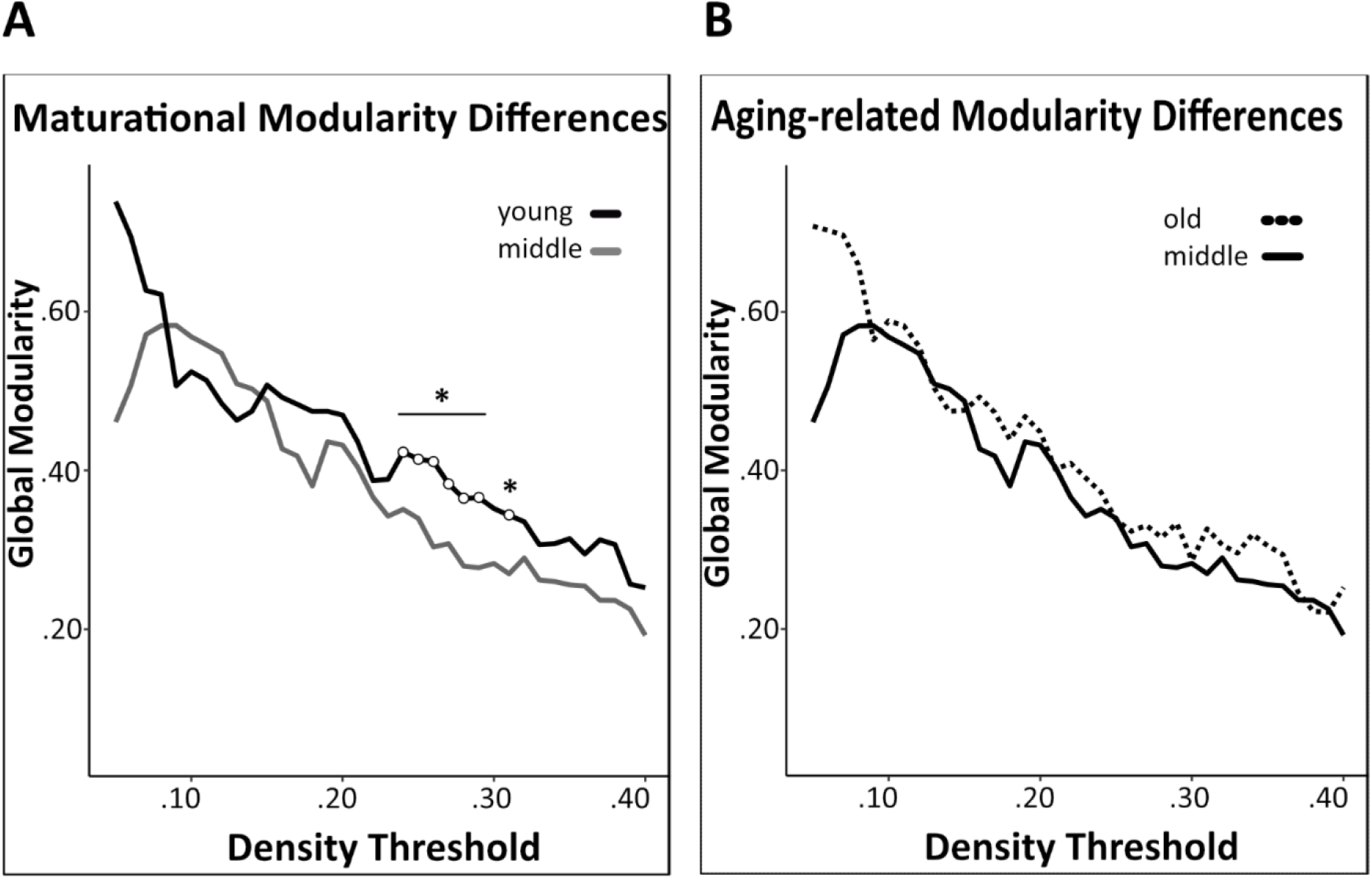
Network modularity comparisons. A significant maturational decrease in modularity was observed between young and middle-age mice networks (**A**). Modularity did not differ significantly between middle-age and old mice networks (**B**). Changes in modularity were determined across density thresholds of 0.05 and 0.40. 5 consecutive significant density thresholds (statistical significance indicated by white circles (*p<0.05)).

Network community modules generated by the Walktrap community detection algorithm revealed densely interconnected subgraphs in the young, middle-age, and old networks (Fig 7A-C). Communities in the young network had densely connected within-community edges (black lines) and few inter-community connections (red lines), while the middle-age and old network communities had progressively fewer within-community connections and more inter-community connections. In the young mice network, the most prominent provincial hubs of their respective communities were the frontal pole, extrapyramidal fiber system, and olfactory areas, while the most prominent connector hubs were the ectorhinal area, temporal association areas. Moreover, in the middle-aged network, the frontal pole, posterior parietal association areas, hypothalamus, pallidum, and lateral forebrain bundle system were identified as provincial hubs, and the temporal association areas, cerebellar nuclei, and pons were identified as connector hubs. Lastly, in the old mice network, the midbrain was identified as the only provincial hub, while the pallidum, prelimbic area, agranular insular areas, and hippocampal formation were identified as connector hubs.

**Fig. 7:**
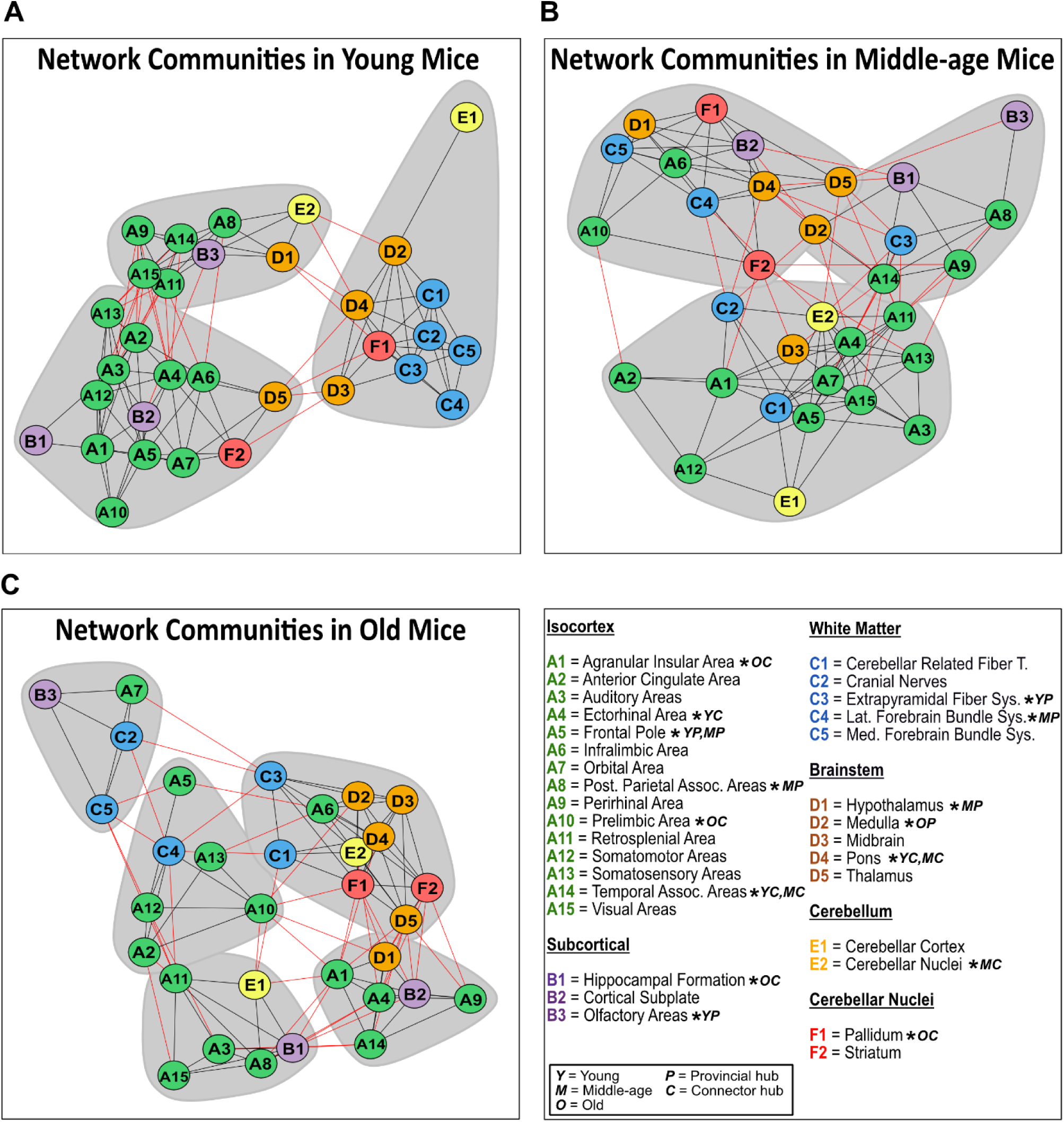
Network communities in mice. Communities of densely interconnected nodes were identified using a Walktrap community detection algorithm for the structural covariance networks of the young (**A**), middle-age (**B**), and old mice (**C**). Regions of interest are sorted by isocortical (green), subcortical (purple), white matter (blue), brainstem (orange), cerebellum (yellow), and cerebellar nuclei (red) regions. Provincial hub regions are denoted by ‘P’, and connector hub regions are denoted by ‘C’ in the young ‘Y’, middle-age ‘M’ and old ‘O’ networks.

## 4. Discussion

Leveraging structural neuroimaging data in young, middle-aged, and old mice, we provide a comprehensive characterization of regional volume and SCN changes across the lifespan, and map them onto working memory performance in the Y-maze paradigm. We observed robust and widespread maturational changes in brain volume, including notable volume loss in isocortical regions accompanied by volume gain in white matter tracts and the brainstem. Aging-related volume changes were more discreet, with further volume loss in select regions of the isocortex, including the orbital area, ectorhinal area, auditory areas, and temporal association areas, coupled with volume gain in white matter tracts, and contrasted by relative preservation in subcortical and cerebellar regions. Additionally, poorer working memory performance on the Y-maze task was associated with smaller volume of the anterior cingulate area and larger volume of the hippocampal formation, regardless of age. Lastly, SCN analyses revealed widespread maturational changes in node-level degree centrality and a significant decrease in network modularity, but not in the aging related comparison. Network transitivity and mean distance remained relatively stable across the mouse lifespan.

### 4.1.1 Brain volume changes across maturation and aging in mice

In the present study, the most prominent maturational changes in volume (young vs. middle-aged mice) occurred in the isocortex. This finding mirrors those in human literature, wherein protracted development in the cerebral cortex precedes widespread gray matter loss, most notably in the prefrontal cortex and to a lesser extent in the parietal and temporal association areas (Raz 1997; Tisserand et al. 2002; Resnick et al. 2003). Similar to these trends, we detected significant volume loss in 7 of the 15 isocortical regions, including 2 of the 3 ‘prefrontal’ regions in rodents (prelimbic area, anterior cingulate area) (Laubach et al. 2018) and the temporal association area. We did not observe significant age-related changes in the remaining prefrontal region (infralimbic area) or in the parietal association area. The aforementioned volume loss in the anterior cingulate area, an *a priori* ROI, aligns with previous rodent literature (Hamezah et al. 2017), along with most human findings (Bergfield et al. 2010; Good et al. 2001; N Raz 1997).

Aging-related volume changes, indexed by comparing middle-age vs. old mice, were also associated with considerable volume loss in the isocortex, with atrophy detected in 4 of the 15 associated regions. Notably, the temporal association area experienced both maturational and aging-related loss, suggesting lifelong volumetric decrease, as did the auditory areas. Moreover, the orbital area, another *a priori* ROI, experienced significant aging-related loss consistent with human literature (Tisserand et al. 2002; Convit et al. 2001), suggesting that this region may be particularly sensitive to late-life volume decrease in mice.

Beyond the isocortex, other areas demonstrated similar patterns to human trajectories of lifespan brain volume. Specifically, we found significant volume increase in 4 of 5 brainstem regions in the maturational comparison. Furthermore, these regions did not experience significant volume loss or gain in the aging-related comparison. These findings align with human trajectories of increasing brainstem volume in maturation (Walhovd et al. 2005), and accounts of relative stability in aging (X. Long et al., 2012; Luft et al., 1999; Walhovd et al., 2011). In addition, the maturational comparison of white matter tracts in the mice revealed 4 of 5 tracts significantly increasing in volume. This aligns with previous rodent studies (Calabrese and Johnson 2013) as well as human trends of white matter gradually maturing into adulthood (Jernigan et al. 2001; Liu et al. 2016).

Although in the present study we described cortical maturational and aging-related volume changes in mice similar to those observed in humans, we also found deviations from human lifespan trajectories. Most notably, we did not detect significant maturational or aging-related volumetric changes in the 3 subcortical structures, including the hippocampal formation, an *a priori* ROI, or in cerebellar regions. This is contrary to evidence from human studies, which suggest that subcortical structures reach maximum volume early in development before undergoing a steep decline in the sixth decade (Dima et al. 2022). It is similarly contrary to evidence of continued hippocampal formation neurogenesis and growth into adulthood in both humans and rodents (E. Sullivan 1995; Diamond, Johnson, and Ingham 1975) and of aging-related volumetric decline of the cerebellum in humans (Jernigan et al. 2001; Naftali Raz et al. 2001; Escalona et al. 1991). Although the absence of aging-related decrease in the hippocampal formation differs from accounts of drastic late-life volume loss in humans and rodents (Jernigan et al. 2001; Zheng et al. 2019; Hamezah et al. 2017; G. Alexander et al. 2012), it does align with contradictory findings of preservation in humans (Good et al. 2001; E. Sullivan 1995). Moreover, it is possible that segmentation of the hippocampal formation may have increased sensitivity to changes in volume (Hackert et al. 2002). Finally, the observed significant aging-related volume increases in 2 of 5 white matter tracts differs from findings of _drastic late-life white matter loss in human literature_ (Jernigan et al. 2001; Liu et al. 2016).

Overall, the volumetric changes found in the present study were more numerous and robust in the maturational comparisons than the aging-related comparisons. We speculate this may be attributable to the relatively shorter late-life period in mice compared to the protracted aging period humans experience. Furthermore, the controlled laboratory setting may have been protective against perturbations caused by adverse environmental variables contributing to aging-related decline. Lastly, it is also possible that our study was underpowered to detect aging-related changes due to small group size and greater inter-subject (i.e., significant within-group) variability.

### 4.1.2 Associations between working memory performance and regional brain volume

Poorer working memory performance in the Y-maze was associated with smaller volume of the anterior cingulate area. Notably, this region also underwent a significant maturational decrease in volume and degree centrality. The anterior cingulate area has previously been implicated in working memory in rodents (Teixeira et al. 2006), similar to the involvement of the ACC in working memory in humans (Kaneda and Osaka 2008), including aging-related decline in functional activity of the ACC in working memory performance (Pardo et al. 2007).

Furthermore, as one of three prefrontal regions in rodents, this association also supports the relationship between the prefrontal cortex and working memory decline (Head et al 2002). In the *a priori* ROI analyses, poorer working memory performance was also associated with larger volume of the hippocampal formation, despite no significant volume differences identified over the lifespan. This association further supports the well-known involvement of the hippocampal formation in working memory in humans, and rodents (Yonelinas 2013; Yoon et al. 2008). A meta-analysis of volumetric changes in the hippocampus and memory outcomes in humans demonstrated high variability in the structural associations in older adults, while youth and young adults trended toward worse performance with higher hippocampus volume (Van Petten 2004). The association observed in our mice is difficult to interpret against the variable human literature, but notably does not align with rodent findings of smaller hippocampal formation volume and poorer working memory performance (Kadar et al. 1990; Hamezah et al. 2017).

We observed additional associations that did not survive multiple corrections. These included the agranular insular area and ectorhinal area, two additional isocortical regions where poorer working memory performance was associated with smaller brain volume. This corroborates the agranular insular area’s involvement in memory processes in rodents, including working memory (DeCoteau et al., 1997; Zhu et al., 2020). The ectorhinal area, which also experienced significant aging-related volume loss, borders the perirhinal cortex and comprises the posterior of the parahippocampal gyrus (Westerhaus & Loewy, 2001). The ectorhinal area is also implicated in memory (Tulving & Markowitsch, 1997), and has been observed to be sensitive to aging-related volume loss in humans (Raz & Rodrigue, 2006b). Lastly, we found that poorer working memory performance was associated with larger volume of the medial forebrain bundle system, a white matter tract implicated in motivation and reward (Anthofer et al., 2015; Wise, 2005). While white matter volume generally shows less associations with cognition compared to those found in gray matter (Taki et al., 2011), it is notable that the significant aging related volume increase in this region deviates from human patterns of white matter loss (Liu et al., 2016). Overall, these associations reflected a general trend of better working memory performance from structurally ‘younger’ brain regions, and declining cognitive performance when regional brain structures were characteristic of older mice.

Although we observed numerous volumetric changes, few brain regions showed associations with working memory performance. This may be due to the relatively discreet significant regional volume changes between middle-age and old mice. It is also possible that older mice utilize cognitive reserve by employing alternative cognitive strategies to compensate for deficits through differential recruitment patterns of brain activity and structural reorganization (Stern 2009; Eyler et al. 2011; Zatorre, Fields, and Johansen-Berg 2012). Lastly, it is notable that aging-related decline in cognitive function is also driven by factors other than brain volume loss, including diminished synaptic, neurotransmitter, cell signaling, and mitochondrial function (Shankar 2010).

### 4.1.3 Structural covariance network topology across maturation and aging in mice

Investigating SCN changes allowed us to consider structural synchronization suggestive of interactions among and between multiple brain regions. The existing SCN literature in rodents is limited (Alexander-Bloch, Giedd, and Bullmore 2013), and to our knowledge no studies have explored lifespan SCN trajectories in mice using whole-brain SCN methodology. In our maturational comparison, 9 of the 32 regions experienced significant changes in degree centrality, with 5 regions in young mice experiencing initially highly connected and influential nodes in the network decrease in centrality into middle-age, including the anterior cingulate area. Four regions emerged as significantly connected nodes in the middle-age network, including the hippocampal formation. These differences in degree centrality largely coincided with significant changes in volume, whereby 3 of the 4 maturational degree centrality decreases co-occurred with significant maturational volume loss, and 3 of the 5 maturational increases in degree centrality accompanied significant maturational volume increase. This may reflect a general trend of higher or lower influence of nodes within the network being associated with directionally consistent volume changes. In contrast, the comparison between middle-age and old mice yielded no significant degree centrality differences across all 32 regions. These findings also offer insight into how mouse SCNs compare to established human lifespan SCN trajectories. In humans, the transition from childhood and adolescence to young adulthood brings a generalized homogenization of covariance patterns, as networks stabilize toward adult topology (Zielinski et al. 2010a). Similarly, a recent large-scale study described pronounced stabilization of overall structural covariance by age 25 (Nadig et al. 2021). The widespread maturational differences in regional degree we observed may reflect developmental reorganization of the SCNs in the young mice, prior to stabilizing into adult SCN patterns in the middle-aged mice. The lack of degree centrality differences between middle-aged and old mice suggest that network stability persisted as mice got older.

In addition to regional connectivity, we investigated global measures of network integration and segregation to further characterize lifespan SCNs in mice. The lack of differences in mean distance across the lifespan of the mice may indicate that mice do not experience late-life reduction in network efficiency in normal aging that is typically observed in human aging (Khundrakpam et al. 2013; Betzel et al. 2014). Furthermore, the lack of maturational or aging relating differences in network transitivity suggests clustering in mouse SCNs is relatively stable across the lifespan. However, we did observe a significant maturational decrease in modularity, whereby young mice had higher network modularity than middle-aged mice, but there were no differences in modularity in the aging-related comparison. Human maturational SCN trajectories demonstrate highly segregated networks favouring localized connections in youth and adolescence, before shifting toward more distributed and globally connected networks in young adulthood (Khundrakpam et al. 2013; Cao et al. 2016). The observed maturational decrease in modularity suggests mice may share a similar trajectory with humans, whereby network segregation decreases from youth to young adulthood. In older age, human SCN segregation has been posited to decrease, becoming further dispersed and disorganized (Chen et al. 2011; Jao et al. 2020), however conflicting findings suggest that network segregation may actually increase in old age (Montembeault et al. 2012). In particular, modularity in structural and functional networks has drawn attention as a potential biomarker for aging (Aboud et al. 2018) as decreasing modularity may be associated with aging-related dedifferentiation of specialized cognitive processes (Chen et al. 2011; Onoda and Yamaguchi 2013; Song et al. 2014). The observed lack of aging-related modularity loss suggests that normal mice aging may differ from SCN segregation trajectories present in human aging. Furthermore, this finding may also reflect the relatively discreet volumetric changes in our aging-related comparison.

To further probe our findings in network modularity, we explored concomitant changes in densely interconnected subgraphs within the networks. The significant maturational decrease in modularity detected in our whole-brain analysis was reflected in the subgraphs, as the high within community connections and low inter-community connections in the young mice communities shifted to sparser within-community and increased inter-community connections in the middle age communities. While the whole-brain aging-related comparison for modularity was not significant, the old-age communities more closely resembled an ‘old network phenotype,’ with few within-community connections, high inter-community connections, and smaller, more numerous communities. Among the identified provincial hubs, the frontal pole emerged as the provincial hub for a community containing predominantly isocortical regions in both young and middle-age networks, suggesting its importance in facilitating specialized information processing among regions associated with higher-order sensory and cognitive processes. Only one provincial hub, the midbrain, emerged in the old-network communities, reflecting the overall reduced within community connections. Lastly, the temporal association areas and pons were identified as connector hubs in both young and middle-aged networks, highlighting their importance in facilitating information exchange between segregated network communities.

Certain limitations in this study should be considered. Most notably, it is possible that that we were underpowered to detect volumetric changes, particularly between the middle-age and old groups, given our relatively small sample size. Future work to validate and further explore regional volume trajectories across the mouse lifespan will benefit from larger sample sizes in each age-group. Secondly, the study utilized a cross-sectional design, considering different groups of mice at each timepoint. Ideally, future studies could employ a longitudinal design, observing the same group of mice over time, thereby increasing the strength of inferences drawn from the maturational and aging-related comparisons. Furthermore, only male mice were available at the time of this study, therefore our results are not generalizable to female mice. Differences in gray matter volume have been observed between male and female mice, both in speed of gray matter development and variation in regional volume (Qiu et al., 2018; Spring et al., 2007). It is possible these inherent differences may influence maturational or aging-related brain volume outcomes. Furthermore, differences in brain structure and function may vary across mice strains. Therefore, the observed changes in brain volume in the Charles River C57BL/6 mice may not be fully generalizable to other strains of mice. Lastly, while working memory decline in aging is well-documented, an additional limitation of our work is that we used only one test, the Y-maze, to evaluate working memory. It is also important to acknowledge that multiple cognitive domains are affected by aging, and future work would benefit from exploring additional domains for a more nuanced picture of the association between volumetric changes and cognitive outcomes.

Despite these limitations, this study provides a detailed account of regional brain volume changes across the mouse lifespan and maps them to working memory performance, while also exploring complementary trends in structural covariance network topology. This comprehensive approach may facilitate future progress in translation between rodent models and human brain aging.

